# 4,5-dihydroxyhexanoic acid is a robust circulating and urine marker of mitochondrial disease and its severity

**DOI:** 10.64898/2026.02.10.705117

**Authors:** Owen S. Skinner, Maria Miranda, Fangcong Dong, Tessa Struhl, Melissa A. Walker, Grigorij Schleifer, Matthew T. Henke, Jon Clardy, Michio Hirano, Darryl C. De Vivo, Eric A. Schon, Kristin Engelstad, Stephanie E. Siegmund, Catherine Laprise, Christine Des Rosiers, Vamsi K. Mootha, Rohit Sharma

## Abstract

Management of patients with mitochondrial respiratory chain diseases is challenging, in part because of our incomplete understanding of pathogenesis and a lack of biomarkers. Unknown metabolites account for >90% of detected features in modern metabolomics experiments and hold immense untapped promise for new basic and biomedical research. We recently used mass spectrometry-based metabolomics to identify and validate 19 circulating blood-based biomarkers for patients with the mitochondrial DNA (mtDNA) m.3243A>G pathogenic variant, which is the most frequent cause of the mitochondrial disorder MELAS (<u>m</u>itochondrial <u>e</u>ncephalomyopathy, lactic <u>a</u>cidosis, and <u>s</u>troke-like episodes). However, the most significantly changing biomarker corresponded to an “unknown” metabolite. Here, we combine cheminformatics with analytical chemistry and identify that feature as 4,5-dihydroxyhexanoic acid (4,5-DHHA), a metabolite previously associated with inherited defects of gamma-aminobutyric acid (GABA) catabolism, but with no prior links to mitochondrial respiratory chain disorders. We validate this finding in an independent MELAS cohort and further show that 4,5-DHHA levels correlate with disease severity and are elevated in patients with other forms of mitochondrial disease and sepsis. Furthermore, brain 4,5-DHHA levels were elevated in two genetic mouse models of mitochondrial disease. *In vitro* and tissue culture experiments indicate that 4,5-DHHA is generated when the GABA catabolite succinic semialdehyde reacts with an intermediate of the pyruvate dehydrogenase reaction and is sensitive to mitochondrial complex I function. Our work identifies 4,5-DHHA as a robust plasma and urine marker of mitochondrial dysfunction in humans and reveals new connections between the respiratory chain and GABA metabolism.

**Significance Statement:** Inborn errors of the mitochondrial respiratory chain cause severe, progressive diseases, yet effective treatments and biomarkers remain limited. Modern metabolomics detects thousands of molecules in biofluids, but the vast majority are unidentified. In this study, we investigate the most significantly altered blood metabolite in patients with the most common mitochondrial disease – MELAS (<u>m</u>itochondrial <u>e</u>ncephalomyopathy, lactic <u>a</u>cidosis, and <u>s</u>troke-like episodes) – and identify it as an 4,5-dihydroxyhexanoic acid (4,5-DHHA). We show that 4,5-DHHA is reproducibly elevated and correlates with severity. Levels are increased across multiple mitochondrial disorders as well as in sepsis and rise when respiratory chain function is impaired. These findings establish 4,5-DHHA as a promising biomarker of mitochondrial dysfunction and reveal a link to dysregulated GABA metabolism.

## Introduction

Respiratory chain diseases (RCDs) arise from genetic lesions to the mitochondrial respiratory chain and cause progressive, often devastating clinical features that can impact any organ system at any age (1, 2). While RCDs are collectively one of the most common inborn errors of metabolism, their clinical management remains challenging and focuses on surveillance and symptomatic management with almost no approved therapies making the treatment of these disorders very challenging (1). Identification of biomarkers for mitochondrial disease can elucidate biochemical mechanisms that may reveal new targetable pathways, provide objective measures to monitor disease progression, and enable grouping of patients based on shared pathomechanisms.

In a recent study, we measured and reported plasma biomarkers from a cohort of patients with MELAS, one of the most common RCDs (3). This Discovery cohort included 102 patients with the m.3243A>G pathogenic variant which accounts for ∼80% of MELAS cases and 32 healthy controls. Twenty of the patients had already suffered stroke-like episodes, which we define here as having MELAS; the remaining 82 are defined as Carriers and together have a spectrum of clinical manifestations. Untargeted metabolomics with liquid chromatography-mass spectrometry (LC-MS) of plasma samples from this cohort revealed classical metabolite markers of mitochondrial disease, like lactate and alanine, that were unambiguously identified using retention time standards. However, numerous unidentified features outperformed identified metabolites and follow-up analytical characterization revealed three families of metabolite biomarkers: lactoyl-amino acids, β-hydroxy acylcarnitines, and β-hydroxy fatty acids. After measuring these markers in an independent validation cohort with m.3243A>G, we validated 19 circulating metabolite biomarkers of MELAS, with many correlating with independent metrics of disease severity. The predominant biochemical signature leading to increases in these validated markers was an elevated NADH/NAD^+^ ratio, termed NADH-reductive stress.

Our prior work underscored the value of mining the unknown “dark metabolome” – which revealed the lactoyl-amino acids and β-hydroxy fatty acids – yet the most significantly changed biomarker in MELAS patients remained unidentified (3). Here, we use a combination of LC-MS/MS and gas chromatography (GC)-MS to identify this metabolite as 4,5-dihydroxyhexanoic acid (4,5-DHHA), which was previously reported to be elevated in patients with a genetic defect of GABA catabolism (4, 5). We find it is also elevated in urine samples from MELAS patients and shows promise as a marker of disease severity. By analyzing our previous human metabolomics studies, we find elevation of plasma 4,5-DHHA also extends to Leigh Syndrome, mitochondrial myopathy, and sepsis, and is stable with exertion. We also demonstrate that 4,5-DHHA is elevated in brain and liver of *Ndufs4*^*-/-*^ and the brains of *Tk2*^*Ki/Ki*^ mouse models of mitochondrial disease and is mechanistically connected to respiratory chain dysfunction.

## Results

### An unknown plasma metabolite marker of MELAS is 4,5-dihydroxyhexanoic acid

To uncover novel, robust markers of disease caused by the m.3243A>G variant, we prioritized unknown metabolites that were significantly elevated in two independent cohorts (3). First, we mined untargeted LC-MS metabolomics of plasma samples from the Discovery cohort. This platform detected and quantified 5584 features, of which we previously identified 237 using retention time standards. An unknown feature with *m/z* 147.0662 and retention time 4.8 min (which we will refer to as U147) showed the most significant (*p =* 1.3 x 10^-10^) separation of MELAS patients and Controls (**Fig. 1A**) among all detected LC-MS features. Notably, U147 was found in all control samples and levels in Carriers were intermediate between Controls and MELAS, with significant pairwise differences among the three groups (**Fig. 1B, left panel**).

**Figure 1.**
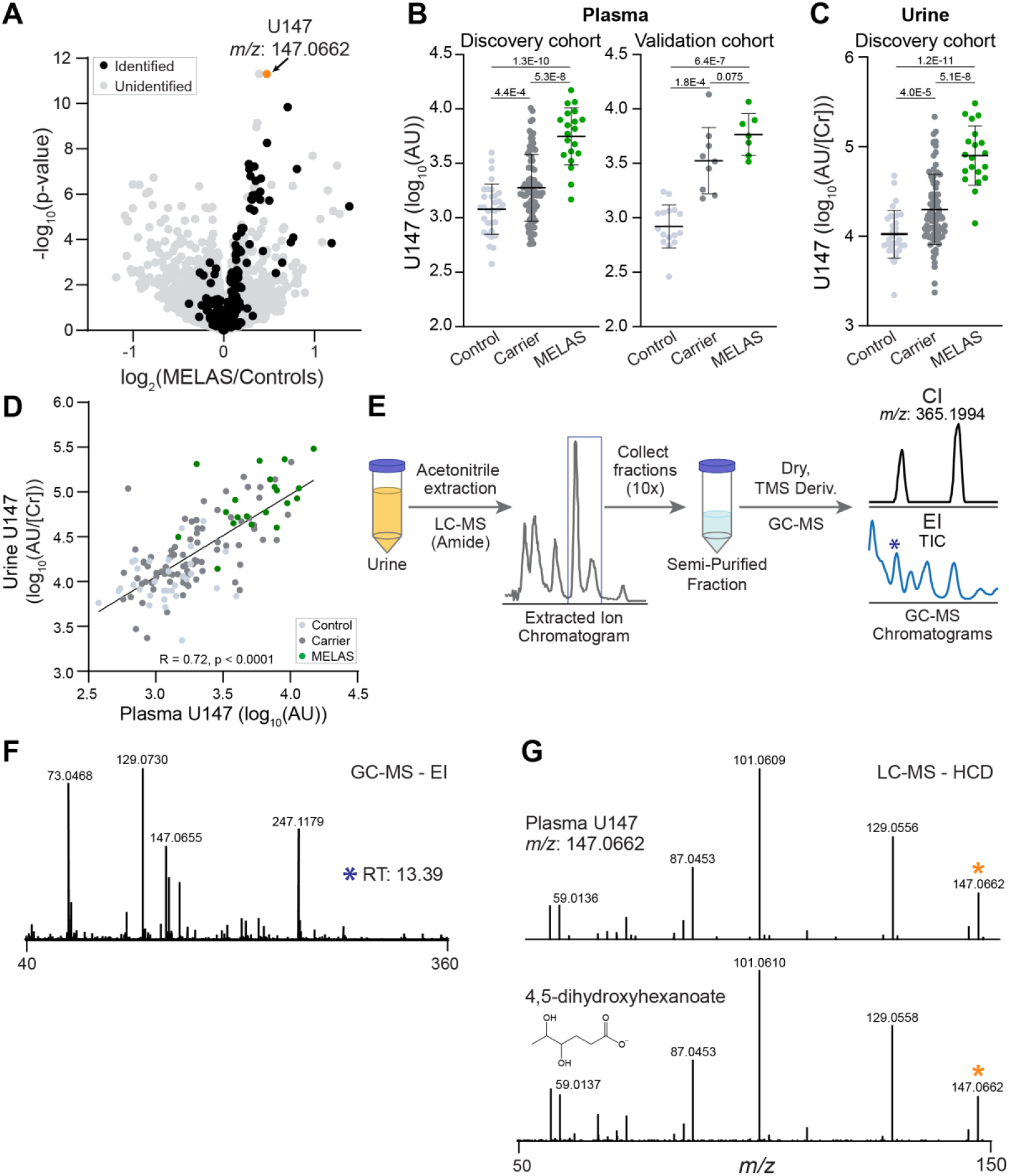
De-orphaning the chemical identity of U147, the top scoring metabolite feature in MELAS. **A)** We mined previously collected untargeted metabolomics data of a Discovery cohort and prioritized 5584 features in the volcano plot (each dot is a feature) by comparing intensities in MELAS patients and Controls. The feature with *m/z* 147.0662 (U147) is one of the strongest markers but its chemical structure is unknown. **B)** U147 is significantly elevated in patients in the Discovery cohort and in an independent Validation cohort. **C)** It is also significantly elevated in urine samples from the Discovery cohort. **D)** U147 levels in plasma and urine are strongly correlated. **E-F)** Chemical extraction, TMS-derivatization and GC-MS with both chemical ionization (CI) and electron ionization (EI) yielded mass and fragmentation patterns that matched published measurements of 4,5-dihydroxyhexanoic acid (4,5-DHHA) (4). **G)** Chemically synthesized 4,5-DHHA standard has the same fragmentation pattern on LC-MS/MS as a peak observed in a sample from a MELAS patient. The precursor ion corresponding to 4,5-DHHA is marked by an orange asterisk.

To validate U147, we looked in an independent Validation cohort of 7 MELAS patients, 9 Carriers, and 16 Controls (collected at different institutes from the Discovery Cohort) and found that U147 was also significantly elevated in Carriers and MELAS patients vs Controls (**Fig. 1B, right panel**). To determine if U147 is also found in urine, we collected LC-MS data on urine samples from the Discovery cohort and found a peak with matching *m/z*, retention time, and fragmentation pattern (**Fig. S1A, B**). The urine level of U147 normalized to urine creatinine concentration (to correct for differences in urine concentration) was significantly elevated in MELAS patients compared to controls, with carriers showing intermediate levels that are significantly different from both MELAS and controls (**Fig. 1C**); importantly, urine U147 strongly outperforms urine lactate in distinguishing the three groups (**Fig. S1C**). Urine and plasma levels of U147 were highly correlated across individuals, suggesting a shared biochemical origin and similar biomarker efficiency (**Fig. 1D**). Together, these data validated U147 as a plasma and urine marker of MELAS and prioritized it for identification and chemical structure elucidation.

The measured *m/z* of U147 was 147.0662 from negative electrospray ionization, which is consistent with the neutral chemical formula C_6_H_12_O_4_, and at least one acidic moiety leading to a [M-H]^-^ ion. We first searched small molecule databases and found multiple compounds with a C_6_H_12_O_4_ chemical formula. We coupled this list with literature searching and compared U147 to pure standards of the cholesterol biosynthesis intermediate mevalonic acid, the coenzyme A precursor pantoic acid, the ω-hydroxylated β-oxidation intermediate 3,5-dihydroxyhexanoic acid, the hydroxylated branched-chain amino acid catabolism intermediate 2,5-dihydroxy-4-methylpentanoic acid, and the GABA catabolite 4,5-DHHA.

Of these candidates, only 4,5-DHHA exhibited a retention time consistent with U147. The first report of 4,5-DHHA came from a prior study of the urine organic acid profiles of patients with succinic semialdehyde dehydrogenase (SSADH) deficiency, an autosomal recessive inborn error of GABA catabolism; this study identified elevations in both the *threo-* and *erythro-* forms of 4,5-DHHA, which has been considered pathognomonic for that condition (4, 6). To determine if the fragmentation of U147 matches the tris-trimethylsilyl(TMS)-derivatized form of 4,5-DHHA reported in the literature, we purified U147 from urine of a healthy donor by collecting fractions from our LC method that contained only U147 at *m/z* 147.0662 and then performed TMS derivatization on this fraction (**Fig. 1E**). Chemical ionization (CI) GC-MS confirmed the intact mass of U147 with the addition of three TMS molecules eluting as two peaks (**Fig. 1E**). Electron ionization (EI) GC-MS with the same published chromatographic method showed fragmentation patterns of both peaks that matched the published pattern of 4,5-DHHA (**Fig. 1F**) (4). Together these results indicate that U147 is indeed 4,5-dihydrohexanoic acid and that the two peaks likely represent the *threo-* and *erythro-* diastereomers (4). To confirm this by LC-MS, we obtained a pure standard of 4,5-DHHA and showed that it has the same HCD fragmentation pattern as U147 in plasma (**Fig. 1G**) and in urine (cosine similarity score of 0.98, **Fig. S1B**), and spiking pure standard into patient plasma and urine samples confirmed that it exhibits the same retention time (**Fig. S1A**). Metabolites that arise from shared metabolic origins often display correlated abundances. We therefore determined the Pearson correlation of 4,5-DHHA with 280 identified plasma metabolites across all samples of the Discovery cohort (**Fig. S1D**) (3). 4,5-DHHA is most strongly correlated with the most changed MELAS markers previously reported but exhibits no particular biochemical patterns and is relatively poorly correlated with plasma lactate (**Fig. S1E**). Together these results demonstrate the successful application of coupled GC-MS and LC-MS methodologies to identify the validated unknown MELAS marker as 4,5-DHHA.

### Levels of 4,5-DHHA are elevated in other mitochondrial diseases and track with severity

We next sought to characterize the clinical context of 4,5-DHHA. Several of the previously reported metabolite markers from these cohorts correlated strongly with metrics of disease severity, indicating their potential as “monitoring biomarkers” (as defined by the US Food and Drug Administration) (3, 7). To determine if 4,5-DHHA could serve as a monitoring marker, we correlated its levels with three severity metrics: urine sediment heteroplasmy, which is the percent of mtDNA in urine sediment carrying the m.3243A>G mutation; Karnofsky performance score, which indicates a person’s ability to perform everyday tasks; and ventricular lactate measured by magnetic resonance spectroscopy which reports the lactate level in the cerebrospinal fluid (CSF) in the ventricles of the brain (3). 4,5-DHHA had one of the strongest correlations with urine sediment heteroplasmy (τ = 0.45) and Karnofsky performance score (τ = -0.41) among all identified metabolites and proteins in our prior study, strongly outperforming lactate (**Fig. 2A, B**). It is also well correlated with ventricular lactate (τ = 0.27) though to a lesser degree than with other identified markers. Creatinine-normalized urine 4,5-DHHA was similarly correlated with urine heteroplasmy, Karnofsky performance scale, and ventricular lactate (**Fig. S2A**), consistent with the high correlation between plasma and urine 4,5-DHHA (**Fig. 1D**). These results suggest that both plasma and urine 4,5-DHHA may serve as monitoring markers of disease severity, although larger longitudinal studies are necessary to rigorously establish how they track over time with severity and specific disease features over time.

**Figure 2.**
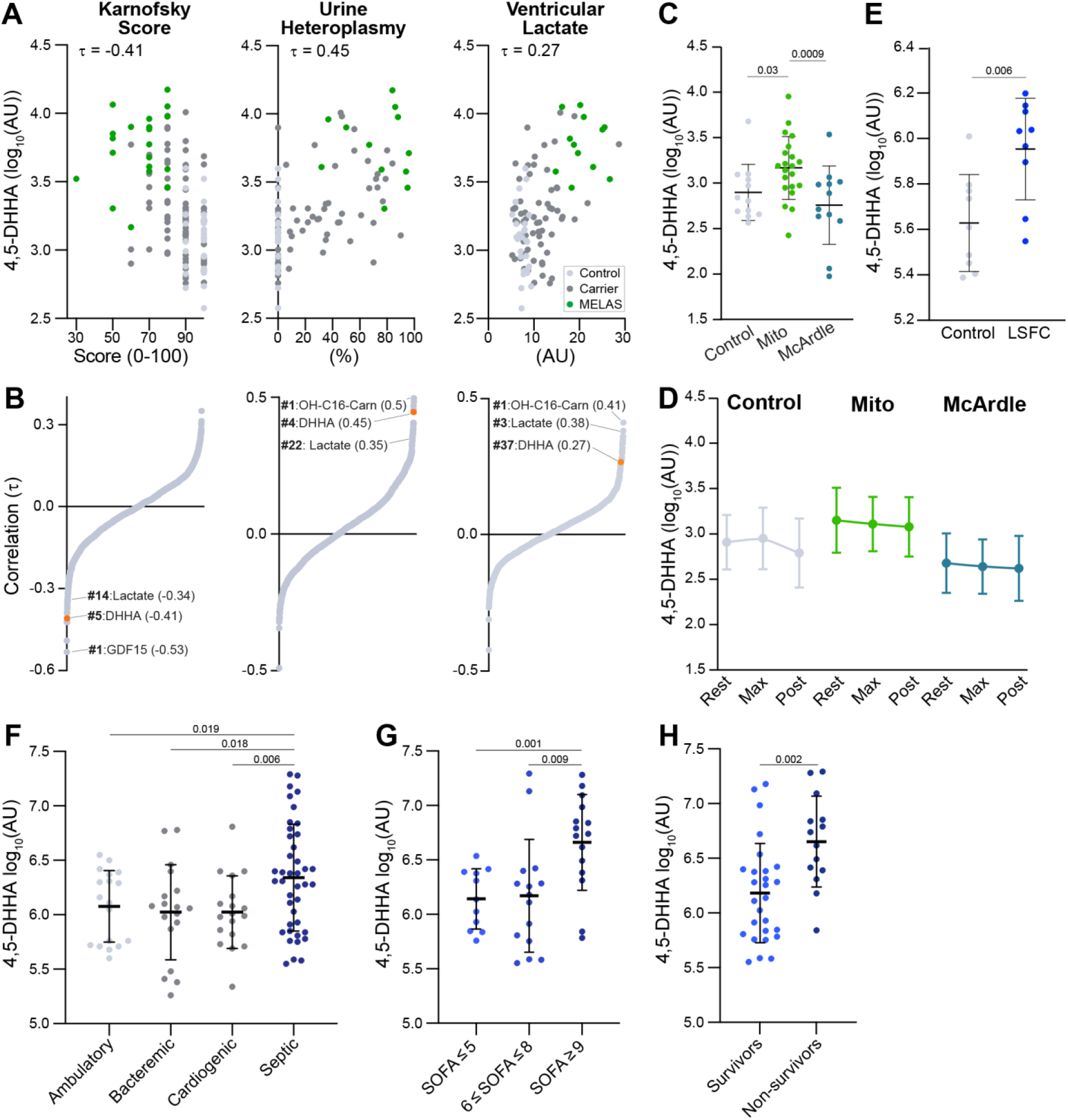
Levels 4,5-DHHA are elevated in multiple patient cohorts and track with severity. **A, B)** Kendall rank correlations of 1973 identified metabolites and proteins with Karnofsky performance score, urine heteroplasmy, and ventricular lactate reveals plasma 4,5-DHHA is one of the strongest markers of severity. **C)** In an exercise metabolomics study of a cohort with healthy controls, mitochondrial myopathy (Mito), and McArdle disease, 4,5-DHHA is significantly elevated in Mito patients and trending lower in McArdle disease. **D)** Plasma 4,5-DHHA is stable through exercise and recovery. **E)** Plasma 4,5-DHHA is elevated in patients with LSFC. **F)** Patients with sepsis have high 4,5-DHHA levels compared to ambulatory controls and patients with bacteremia or cardiogenic shock. **G-H)** Among patients with sepsis, plasma levels of 4,5-DHHA are higher in those with Sequential Organ Failure Assessment (SOFA) score > 9 (greater severity) and in non-survivors.

Biomarkers can also be used to diagnose and categorize diseases (8). The elevated levels of urine 4,5-DHHA in SSADH deficiency patients and now in plasma and urine in MELAS patients already indicate its disease specificity is broader than previously appreciated. Considering the shared biochemical origin of previously described RCD markers, we wondered if circulating 4,5-DHHA is elevated in patients with other mitochondrial diseases. To address this, we mined two untargeted metabolomics datasets associated with previously reported studies of patients with mitochondrial dysfunction. First, in a prior study of mitochondrial myopathies resulting from a range of mtDNA genetic lesions, we compared the metabolic responses to exercise in three groups: 1) healthy controls (N=12), 2) patients with mitochondrial myopathies (N=21), and 3) patients with McArdle disease (N=12). McArdle disease is an autosomal recessive condition due to lesions in the muscle isoform of glycogen phosphorylase (*PYGM*) which prevents mobilization of glycogen stores (9). All participants performed an exercise cycle challenge until maximal exertion, and blood samples were collected at rest, peak exercise, and 10 minutes post exercise. At rest, plasma 4,5-DHHA was significantly elevated in mitochondrial myopathy compared to controls and McArdle disease (**Fig. 2C**); levels in McArdle patients trended lower than controls but fell short of meeting statistical significance. Importantly, 4,5-DHHA levels remained stable through exercise, in contrast to the classical RCD biomarkers lactate or pyruvate or the recently discovered lactoyl-amino acids (**Figs. 2D, S2B-C**) (3, 10).

Second, we collected LC-MS data using the same method on plasma samples from a prior study of Leigh Syndrome French Canadian Variant (LSFC). LSFC is an autosomal recessive RCD caused by pathogenic variants in the nuclear gene *LRPPRC* which encodes a protein necessary for translation of subunits of complex IV (11, 12). This study consisted of plasma samples collected from 9 patients with LSFC and 9 healthy controls who were matched for age, sex, body mass index (BMI), and activity level. In these patients, we again found that plasma 4,5-DHHA was significantly elevated in mitochondrial disease (**Fig. 2E**). Together, these results show that elevated 4,5-DHHA extends beyond SSADH deficiency and MELAS to include multiple mitochondrial diseases caused by mtDNA and nuclear DNA lesions, correlates with disease severity, and importantly, is not sensitive to exertion-induced variability which may improve its clinical reliability over other markers.

### 4,5-DHHA is also elevated in sepsis and predicts mortality

We next wondered if more common conditions associated with secondary respiratory chain dysfunction might also show elevated 4,5-DHHA. Since sepsis has recently been demonstrated to share metabolic markers with RCDs, we mined a recently collected dataset that included patients with septic shock (N=42), cardiogenic shock (N=19), bacteremia without sepsis (N=18), and ambulatory controls (N=19) (13). These measurements show that 4,5-DHHA is higher in patients with septic shock compared to each of the three other cohorts (**Fig. 2F**). Among patients with sepsis, 4,5-DHHA levels were highest in the subsets with the highest Sequential Organ Failure Assessment (SOFA) score and among non-survivors (**Fig. 2G, H**). Together, these results demonstrate that circulating 4,5-DHHA is a strong marker of sepsis and correlates with severity in this disease.

### Levels of 4,5-DHHA are markedly elevated in the brains of mouse models of mitochondrial disease

Studies of 4,5-DHHA in patients with SSADH deficiency have found its levels to be elevated in urine, brain, cerebrospinal fluid, liver, and kidney (4–6). To delineate the tissue distribution of 4,5-DHHA in RCD, we turned to the *Ndufs4*^*-/-*^ mouse model of Leigh Syndrome, which closely aligns with human disease features (14). Due to a deficiency in a subunit of complex I of the respiratory chain, *Ndufs4*^*-/-*^ mice develop weight loss at day ∼30 followed by progressive neurological decline manifesting as seizures, ataxia, hypothermia, with a median lifespan of approximately 55 days. We measured 4,5-DHHA levels in liver, heart, kidney, quadriceps, and brain in 56-day-old WT and *Ndufs4*^*-/-*^ mice (n=4 male, 4 female) but were unable to detect it in mouse plasma. At this advanced point of disease progression, we found 4,5-DHHA was significantly elevated in liver (**Fig. 3A**) and markedly so in brain (**Fig. 3B, left side**). Since hypoxia therapy has been found to dramatically reverse many disease features even in mice with late-stage disease, we sought to determine the effect of hypoxia on brain levels of 4,5-DHHA (15, 16). In mice raised for 55 days at normoxia and then transitioned to 11% hypoxia for 42 days (i.e., age 98 days), we found that while disease features improved as previously reported, 4,5-DHHA levels remained elevated (**Fig 3B, right side**).

**Figure 3.**
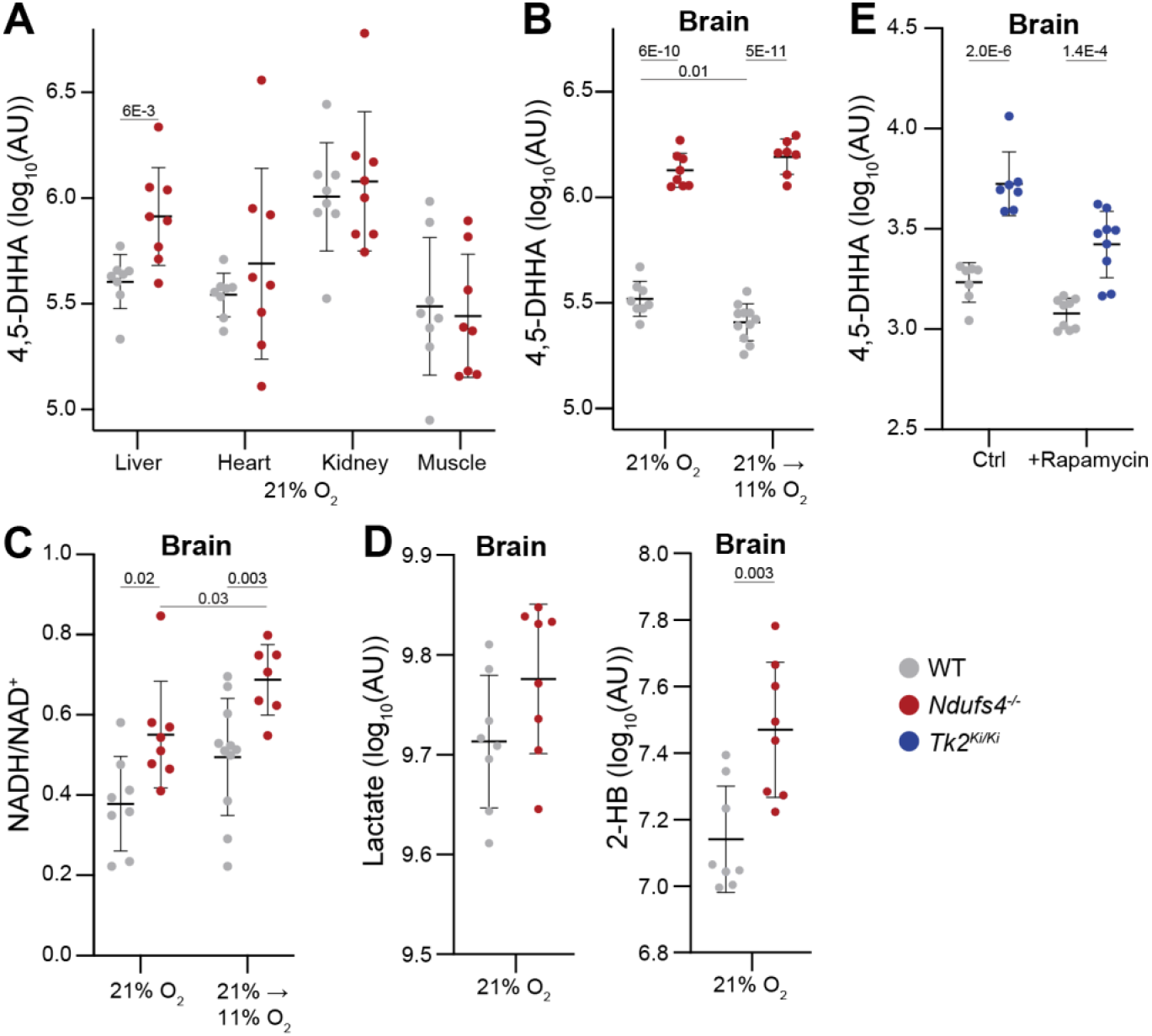
4,5-DHHA is elevated in two genetically distinct mouse models of mitochondrial disease. **A)** In 56-day old mice reared at 21% oxygen, 4,5-DHHA is elevated in liver of *Ndufs4*^*-/-*^ mice with advanced disease. **B)** The 4,5-DHHA level in brain is elevated in 56-day old mice reared at 21% oxygen (left side) and remains elevated after mice are transitioned to 11% oxygen for 42 days rescuing symptoms (98-days old, right side). **C)** The relative NADH/NAD^+^ ratio in brain is elevated in 56-day old mice reared at 21% oxygen (left side) and remains elevated after mice are transitioned to 11% oxygen for 42 days (98-days old, right side). **D)** In 56-day old mice reared at 21%, the brain level of lactate is unchanged in *Ndufs4*^*-/-*^ mice and 2-hydroxybutyrate (2-HB) is elevated. **E)** In *TK2*^*Ki/Ki*^ mouse brain, 4,5-DHHA level is markedly elevated at 15 days and is partially rescued by rapamycin treatment.

GABA catabolism proceeds through two steps in the mitochondrial matrix: a transamination reaction catalyzed by GABA-transaminase (GABA-T, gene *ABAT*) forming succinate semialdehyde, followed by an NAD^+^-requiring oxidation reaction catalyzed by SSADH (gene *ALDH5A1*) (17, 18); both enzymes are located in mitochondria (19–21). In light of this requirement for NAD^+^ and our prior observation of the prominent role of NADH-reductive stress in mitochondrial disease, we measured NADH/NAD^+^ ratios in brain tissues from these mice. Indeed, the relative NADH/NAD^+^ in brain was elevated in 56-day old *Ndufs4*^*-/-*^ mice reared in 21% oxygen and was not corrected after 42 days of 11% hypoxia (**Fig. 3C**). We compared the elevation of 4,5-DHHA in brain with two other metabolite markers of the *Ndufs4*^*-/-*^ mouse model – lactate and 2-hydroxybutyrate (**Fig. 3D**) – and found that only the latter is significantly elevated in brain, though not to the extent of 4,5-DHHA (15, 16). Together, these results reveal that while 4,5-DHHA is found in all tissues measured in the *Ndufs4*^*-/-*^ mouse model, it is elevated only in brain and liver and brain 4,5-DHHA levels and NADH/NAD^+^ do not respond to hypoxia rescue therapy.

To determine if elevation of 4,5-DHHA in brain also extends to another RCD, we re-analyzed previously collected data from a knock-in mouse model of thymidine kinase 2 (TK2) deficiency (*Tk2*^*KI/KI*^, referred to here as the TK2 mouse) which impairs mitochondrial nucleotide salvage and leads to progressive mitochondrial DNA depletion and respiratory chain disease (22). Newborn TK2 mice initially resemble WT mice but after approximately one week develop rapidly progressive failure to thrive and motor dysfunction, with death occurring at a median of two weeks. Previous work showed low-dose rapamycin extends lifespan by approximately 10 days (22). Brain measurements in 15-day-old mice showed markedly elevated 4,5-DHHA which partially responded to rapamycin treatment (**Fig. 3E**). While metabolic profiling was also performed on liver, we did not obtain a clear LC-MS peak for 4,5-DHHA and NADH and NAD^+^ were not measured in these tissues. In sum, these results show brain levels of 4,5-DHHA are elevated in a mouse model of TK2 deficiency and partially respond to rapamycin therapy.

### Biochemical origin of 4,5-DHHA

As noted above, 4,5-DHHA was first identified in urine of patients with SSADH deficiency, an inherited defect of GABA catabolism which occurs in the mitochondrial matrix. After GABA is transported into mitochondria through a yet unidentified transporter, it is first transaminated by GABA aminotransferase to succinic semialdehyde (SSA) (23, 24). In the final step, succinic semialdehyde dehydrogenase (SSADH) oxidizes SSA to the TCA cycle intermediate succinate using NAD^+^. Previous work hypothesized that the formation of 4,5-DHHA begins with a spontaneous condensation between SSA and an intermediate of the pyruvate dehydrogenase complex (PDC) reaction (**Fig. 4A**) (4, 25).

**Figure 4.**
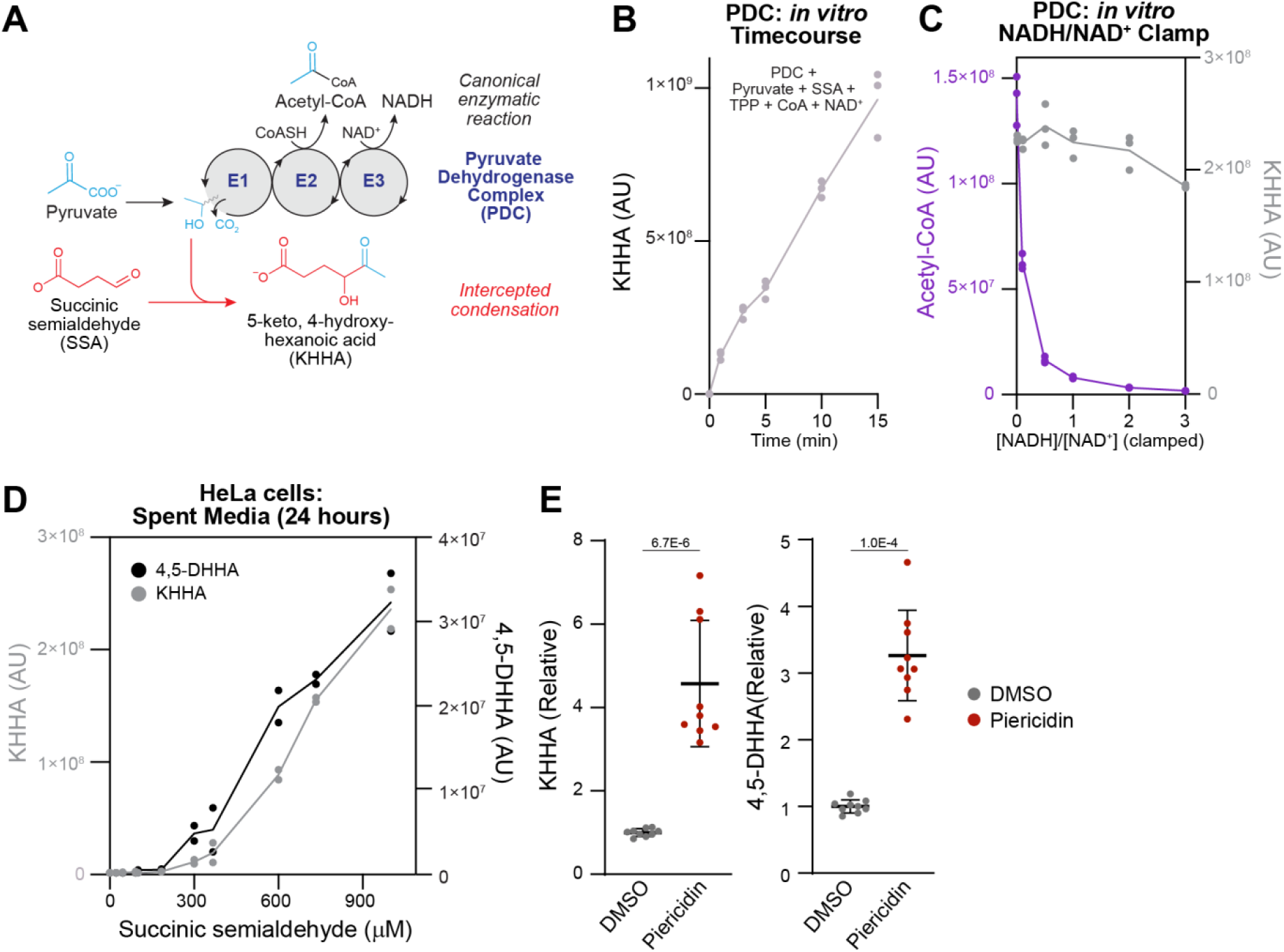
Respiratory chain dysfunction and pyruvate availability contribute to elevated 4,5-DHHA. **A)** In the canonical PDC reaction, the E1 component decarboxylates pyruvate leaving an activated two-carbon intermediate which is released as acetyl-CoA by E2 and two electrons are captured in NADH by E3. The first step in the proposed pathway for 4,5-DHHA synthesis is a condensation between SSA and the two-carbon intermediate forming KHHA. **B)** Incubation of pyruvate and SSA with PDC (with thiamine pyrophosphate (TPP), coenzyme A (CoA), and NAD^+^) produces KHHA *in vitro*. **C)** When this reaction is performed at a range of NADH/NAD^+^ ratios, acetyl-CoA production declines as the ratio increases but KHHA production is relatively unchanged. **D)** Measurements of spent media after HeLa cells are incubated with a range of SSA concentrations for 24 hours show production of both KHHA and 4,5-DHHA. **E)** Treatment of HeLa cells with piericidin (in the presence of 500 μM SSA) increases KHHA and 4,5-DHHA production. Data points are relative to the means of KHHA and 4,5-DHHA in the presence of DMSO.

The canonical PDC reaction sequence begins with the E1 component decarboxylating pyruvate, leaving the “activated” two-carbon hydroxyethyl intermediate bound to thiamine pyrophosphate (26). Next, the E2 component binds the intermediate through its lipoate cofactor and releases it as acetyl-CoA, leaving lipoate in the reduced form. Finally, lipoate is re-oxidized by the E3 component using its FAD cofactor to transfer two electrons to NAD^+^, forming NADH. Prior work indicates that SSA can intercept and condense with the hydroxyethyl intermediate bound to E1, forming 5-keto-4-hydroxyhexanoic acid (KHHA) (25). To confirm that this mechanism can generate KHHA *in vitro*, we first measured the condensation reaction between SSA and the two-carbon intermediate of PDC. We incubated SSA and pyruvate with porcine PDC in the presence of pyruvate, thiamine pyrophosphate (the cofactor for the E1 component), NAD^+^ and coenzyme A. This reaction showed steady production of KHHA over time (**Fig. 4B**), but no detectable 4,5-DHHA, providing validation that PDC catabolism of pyruvate in the presence of SSA is sufficient to generate KHHA but not 4,5-DHHA. KHHA is thought to be subsequently reduced to 4,5-DHHA by an unknown dehydrogenase (4). Notably, the spontaneous condensation reaction forming KHHA is expected to produce both (*R-*) and (*S-*) hydroxyl enantiomers at the 4-position, while the dehydrogenase converting KHHA to 4,5-DHHA likely generates a single hydroxyl enantiomer at the 5-position. The combination of enantiomers at these two chiral carbons explains the observation of the *threo-* and *erythro-* diastereomers detected in our purification (**Fig. 1E**).

To further interrogate the biochemical mechanism connecting mitochondrial dysfunction with 4,5-DHHA accumulation, we first wondered if *in vitro* KHHA formation is sensitive to the NADH/NAD^+^ ratio in the context of complete PDC turnover. We repeated the *in vitro* KHHA assay in the presence of coenzyme A, NAD^+^ and NADH clamped at six different NADH/NAD^+^ ratios and measured acetyl-CoA and KHHA production after five minutes. Consistent with the dependence of PDC on NAD^+^ and its well-established inhibition by high NADH, acetyl-CoA production was markedly decreased by increasing NADH/NAD^+^ (**Fig. 4C**). However, KHHA production remained largely unchanged through a range of NADH/NAD^+^ ratios (**Fig. 4C**) indicating that increased NADH/NAD^+^ is not sufficient to cause elevated KHHA formation from PDC. Again, 4,5-DHHA was not generated at detectable levels during this experiment. This suggests that while NADH-reductive stress inhibits PDC activity to produce acetyl-CoA, in the presence of SSA, E1 activity can surprisingly continue – uncoupled from E2 and E3 – to allow nonspecific condensation with SSA to produce KHHA.

Next, to test the role of mitochondrial respiratory chain dysfunction in 4,5-DHHA production, we turned to HeLa cells which express SSADH at low levels. When we added SSA to HeLa cells in increasing amounts, we observed a corresponding increase in media levels of both KHHA and 4,5-DHHA after 24 hours (**Fig. 4D**). We selected an intermediate concentration of 500 μM SSA for further experiments. Addition of the complex I inhibitor piericidin caused a significant increase of both KHHA and 4,5-DHHA production (**Fig. 4E**). In sum, these results indicate that SSA addition to cultured cells is sufficient to drive 4,5-DHHA production that is increased with chemical inhibition of complex I, which aligns with our results in *Ndufs4*^*-/-*^ mice and human patients (**Fig. 3B**).

While these experiments do not demonstrate inhibition of SSADH induced by respiratory chain dysfunction, they support a mechanism in which accumulation of SSA drives condensation with the hydroxyethyl intermediate of the E1 component of PDH forming KHHA (**Fig. 5**). The enzyme that converts KHHA to 4,5-DHHA remains unknown but likely involves a reduction reaction which would be favored in the setting of an elevated NADH/NAD^+^ ratio.

**Figure 5.**
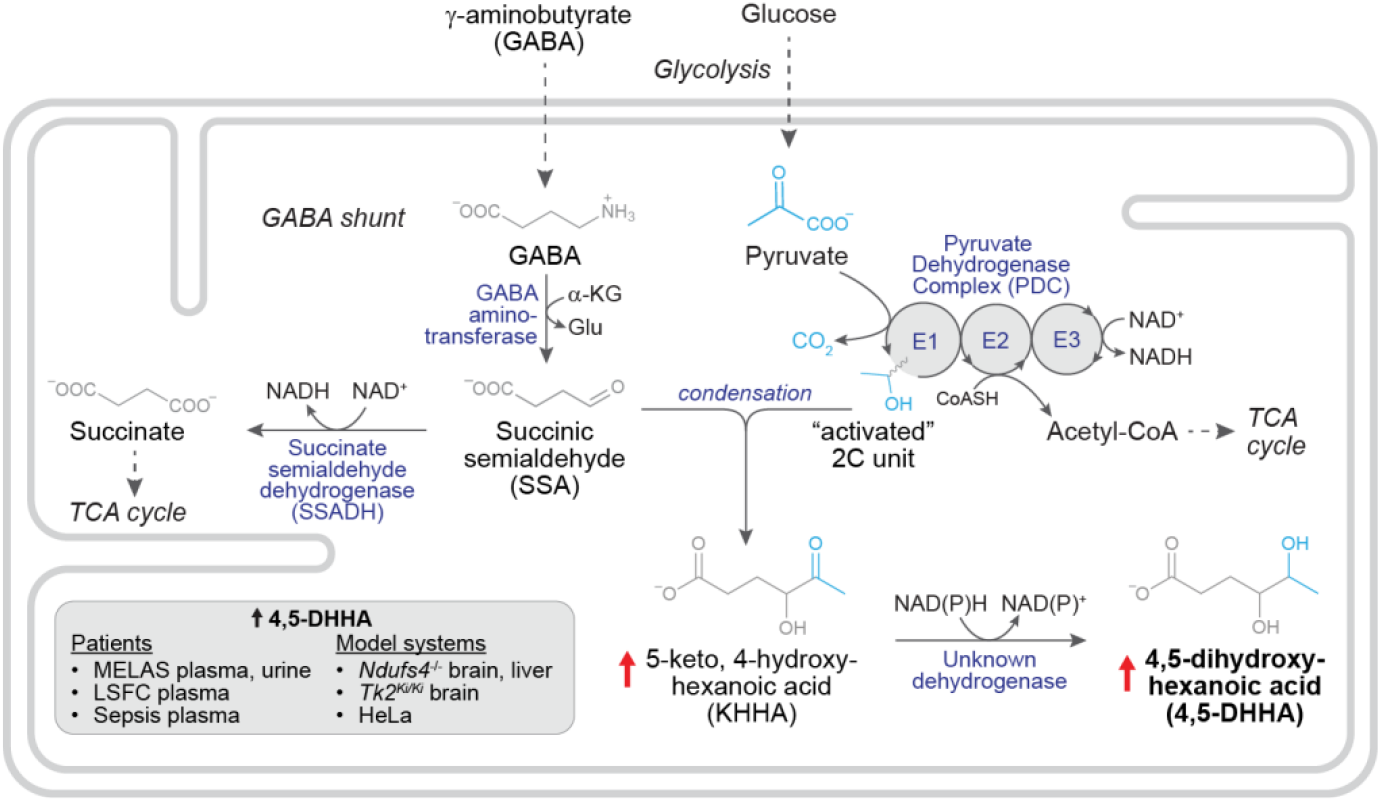
Proposed mechanism leading to 4,5-DHHA elevation with respiratory chain dysfunction. Respiratory chain inhibition from inherited lesions or chemical inhibition inhibits SSADH leading to elevation of SSA which condenses with the “activated” two carbon intermediate of the E1 component of PDC producing KHHA. Reduction of KHHA to 4,5-DHHA by an unknown enzyme would presumably be favored in the setting of NADH-reductive stress; note that the identity and compartmental location of this enzyme is unknown.

## Discussion

Unknown metabolites hold immense untapped promise for new insights into the biochemical fingerprints of diseases and novel mechanistic processes. However, chemical identification is challenging, and prioritizing features to pursue can be laborious. Here, we focused on an unknown metabolite that was one of the strongest plasma markers in two previously studied independent MELAS cohorts, which provided strong motivation for its identification. A combination of analytical measurements, cheminformatics, and literature revealed it to be 4,5-DHHA – a metabolite previously linked to SSADH deficiency. By analyzing prior studies, we highlight 4,5-DHHA as a potential “monitoring biomarker” of MELAS severity and find it is also elevated in other mitochondrial diseases as well as in sepsis. *In vitro* and tissue culture models confirm that 4,5-DHHA elevation is promoted by respiratory chain inhibition. The patient measurements, mouse models, and biochemical studies together provide insights into the biochemistry of 4,5-DHHA, pathophysiology of RCDs, and translational application of this new biomarker.

### Biochemical insights

Our discovery of 4,5-DHHA as an RCD biomarker suggests an unexpected, shared metabolic root process with SSADH deficiency, and our *in vitro* and tissue culture experiments provide mechanistic insights into its elevation with respiratory chain dysfunction. We propose that 4,5-DHHA elevation with RC inhibition is driven by a three-step process: 1) secondary inhibition of SSADH leads to accumulation of SSA, 2) SSA condenses with the hydroxyethyl intermediate of the E1 component of PDC forming KHHA, and 3) KHHA is reduced to 4,5-DHHA by an unknown dehydrogenase (**Fig. 5**). Notably, we do not detect KHHA in human plasma, suggesting that this third step is highly efficient in humans or that 4,5-DHHA can be more efficiently released from cells.

We speculate that levels of the two substrates for 4,5-DHHA synthesis – SSA and pyruvate – influence its circulating levels. While our HeLa cell experiments demonstrate complex I inhibition leads to elevation of 4,5-DHHA, this system expresses only SSADH and lacks GABA transaminase and is therefore unable to reveal how respiratory chain inhibition alters SSA levels *in vivo*. Respiratory chain inhibition may induce secondary inhibition of SSADH activity through at least two mechanisms. First, NADH-reductive stress caused by respiratory chain dysfunction may inhibit SSADH, since elevated NADH has been noted to strongly inhibit this enzyme (Ki of ∼100 μM) (27–29). Second, the catalytic loop of SSADH has been shown to be a redox switch that is sensitive to reactive oxygen species (i.e., oxidizing stress) and can form a disulfide bond, inhibiting activity. Additional experiments will be necessary to demonstrate elevation of SSA and dissect this mechanism.

Our *in vitro* studies of the second step imply that in the presence of SSA a fraction of the PDC E1 intermediate are intercepted in a condensation with SSA at a rate that does not change even when high NADH/NAD^+^ inhibits acetyl-CoA formation by E2. These measurements provide a clear demonstration of this reaction for 4,5-DHHA synthesis and have biochemical and physiological implications. First, the constant rate of KHHA production through a range of NADH/NAD^+^ ratios suggests SSA access to the hydroxyethyl intermediate of E1 is constrained under these conditions, which may be explained by sequestration of the intermediate in the E1 active site. Second, it raises the possibility of a family of such condensation products arising from reactions between aldehydes and intermediates of three-component α-keto acid dehydrogenases. In fact, such products have been reported in classic and recent studies (25, 30–33) and have even been proposed as biomarkers, including the report of 4,5-dihydroxyheptanoic acid which may arise from a similar process between SSA and 2-ketobutyrate (6, 34). Finally, this process reveals a potential source of energy loss in the PDC reaction which appears to occur even in healthy individuals, as we observed 4,5-DHHA even in this cohort. Future work can explore whether 4,5-DHHA is biologically active, catabolized, a dead-end metabolite, or if it can be reclaimed by a proof-reading enzyme.

Our measurements of circulating 4,5-DHHA and lactate in McArdle deficiency provides insight into the systemic regulation of this metabolite. Due to a lack of muscle glycogen phosphorylase, these patients cannot mobilize muscle glycogen and therefore do not mount a lactate response to exercise (**Figs. S2B**). While the role of the GABA shunt in the liver is not fully understood, the most parsimonious explanation for lower circulating levels of 4,5-DHHA in McArdle deficiency (**Fig. 2C**) is decreased lactate flux to the liver via the Cori cycle resulting in a diminished hepatic pyruvate pool and reduced 4,5-DHHA production.

### Pathophysiological insights

An emerging view of the biochemical consequences of RCDs is that disrupted respiratory chain activity triggers proximal metabolic changes like ATP deficiency, elevated NADH/NAD^+^, unused oxygen, or ROS formation. These in turn disrupt multiple biochemical pathways, unleashing numerous distal metabolic perturbations including lactate, alanine, lactoyl-amino acids, and now, 4,5-DHHA each of which may contribute to or buffer pathogenesis or may simply be inert. Connecting biochemical ripples through mechanistic steps to disease features can help unravel pathogenesis, reveal therapeutic approaches, and clarify the translational scope of biomarkers.

Our work provides a framework for this process through two promising therapies. We find that while hypoxia markedly rescues many of the neurological symptoms in *Ndufs4*^*-/-*^ mice, NADH-reductive stress and 4,5-DHHA are not ameliorated in whole brain from these animals. Importantly, our work cannot reveal differences of these metabolites within critical regions of the brain impacted in these mice; additionally, we did not undertake full functional characterization of these mice and cannot correlate changes to specific neurological symptoms. Suppressor genetics studies in *C. elegans* recently indicate that hypoxia helps to preserve partial forward electron flow in mutant complex I to help to defend the proton motive force (PMF) (35). While that small amount of forward electron flow in *C. elegans* is sufficient to ameliorate many aspects of the neurological disease, it seems that the brain reductive stress in *Ndufs4*^*-/-*^ mice is not fully reversed. On the other hand, the mTOR inhibitor rapamycin extends lifespan in the TK2 mouse and causes partial rescue of 4,5-DHHA. Together, these observations underscore the multiple pathways that likely contribute to RCD pathogenesis and the importance of mechanistic clarity and detailed symptom correlation in biomarker evaluation.

The discovery of 4,5-DHHA highlights the potential role of disrupted GABA metabolism in RCDs and has important mechanistic and therapeutic implications. While we were unable to measure GABA levels, it has previously been reported to be lower in *Ndufs4*^*-/-*^ brain, which is restored by rapamycin (36). Combined with our observations, this raises the possibility that diminished GABA levels may be partly due to shunting towards 4,5-DHHA. The significant neurological features seen in inherited lesions of GABA catabolism raises the possibility of shared pathophysiological processes with RCDs which warrant further investigation (4). Patients with SSADH deficiency have a relatively static presentation of intellectual disability, seizure disorders, and behavioral issues, but are not known to suffer acute decompensation in the face of metabolic stress – a significant threat for MELAS patients (37, 38). Finally, experience from SSADH deficiency management indicates utilization of GABA-modulating anti-seizure medications targeting this pathway (e.g. tiagabine, vigabatrin) should be approached with caution (37).

### Translational potential for 4,5-DHHA

Our discovery of 4,5-DHHA as a robust marker of MELAS and other RCDs brings the metabolic profile of these conditions into greater focus and our results provide insights into its specificity and sensitivity. Elevation of 4,5-DHHA in MELAS, mitochondrial myopathies, LSFC, as well as sepsis indicate the importance of considering the clinical scenario as well as the complete metabolic profile of the patient. In contrast to lactate and lactoyl-amino acids, the stability of 4,5-DHHA during exertion may improve its clinical sensitivity and specificity.

The role of pyruvate as a substrate in 4,5-DHHA synthesis is underscored by the lower levels observed in McArdle disease and predicts lower circulating levels of this metabolite will be a feature shared among glycogen storage diseases impacting glycogen mobilization. Additionally, we speculate that given the central role of PDC in endogenous 4,5-DHHA synthesis, patients with PDC defects disrupting the decarboxylase activity in the E1 component (due to inherited lesions or thiamine deficiency) will also have lower 4,5-DHHA levels. We note that recent work demonstrated that in a mouse model of sepsis thiamine deficiency inhibits PDC activity and underlies hyperlactatemia; our measurements suggests that our cohort of patients with sepsis retained sufficient E1 activity to produce 4,5-DHHA (39). Finally, though we did not assess the effects of fasting and feeding on 4,5-DHHA, our results suggest plasma levels will rise in the fed state when pyruvate availability increases.

Taken together, our work highlights analytical and cheminformatic strategies to successfully prioritize, identify, and characterize an unknown metabolite from the dark metabolome. Here, we spotlight 4,5-DHHA – a previously little-known metabolite only associated with a defect of GABA catabolism – as a plasma and urine metabolite that is robustly elevated in RCDs and sepsis. 4,5-DHHA shows promise as both a diagnostic and monitoring biomarker and is correlated with metrics of disease severity in both MELAS and sepsis and is stable to exertion. Our observation of 4,5-DHHA elevation in two mouse models further extends its translational applicability and highlights its marked elevation in the brain. *In vitro* and cell culture experiments point to a mechanism involving PDC, the GABA shunt intermediate SSA, and complex I inhibition. In sum, this work reports 4,5-DHHA as a robust biomarker of multiple forms of respiratory chain deficiency and may serve as a critical component in the biochemical fingerprints of these conditions.

### Limitations

Our work provides a foundation for the clinical application of 4,5-DHHA but has important biochemical and translational limitations to be addressed in future studies. An additional reaction that could, in theory, produce KHHA is a similar condensation reaction between acetaldehyde and the succinyl-thiamine pyrophosphate intermediate of the E1 component of α-ketoglutarate dehydrogenase. A key mechanistic piece that remains unknown is the dehydrogenase that reduces KHHA to 4,5-DHHA. *In vitro* tests of malate dehydrogenase and lactate dehydrogenase indicated these enzymes were unable to perform this reaction (data not shown). We confirmed the identity of 4,5-DHHA with a combination of retention time and MS/MS fragmentation for the Discovery cohort (plasma and urine), Validation cohort, sepsis cohort, and *in vitro* and cell culture experiments; the identity was based on retention time alone in the exercise, LSFC, and mouse studies. Of note, we were unable to unambiguously detect KHHA in the mouse and human samples and we did not determine the enantiomeric form of 4,5-DHHA in our samples. Finally, while urine organic acid measurements have detected 4,5-DHHA, to our knowledge current clinical assays are incapable of measuring plasma levels, a hurdle facing its clinical implementation.

## Materials and Methods

### Study cohorts and study design

All samples and clinical information were collected following approval of a secondary use protocol by the Institutional Review Boards at Massachusetts General Hospital. The **Discovery** and **Validation cohorts** which were first utilized and described in detail in a prior study (3). Briefly, the Discovery Cohort consists of 82 Carriers, 20 MELAS patients (all of whom carry the m.3243A>G variant) and 32 controls. For the Discovery Cohort, blood and urine samples were collected in the morning and magnetic resonance spectroscopy (MRS) was performed within two days. Among the 82 Carriers, one did not provide a urine sample and two additional Carriers provided urine samples but no plasma sample; thus, there are 83 urine samples from Carriers and 81 of these have paired plasma samples. Ten individuals with the m.3243A>G variant required food with their morning medications which was taken >2 hours prior to sample collection; all other participants were fasting at the time of sample collection. MRS was performed on 27 controls, 65 Carriers, and 12 MELAS patients. Urine heteroplasmy, MRS lactate, and Karnofsky scores of the Discovery cohort were determined as previously described (3). The Validation cohort consisted of 9 Carriers, 9 MELAS patients (all with the m.3243A>G variant), and 9 controls. The **Exercise cohort** has been described in detail previously consisted of 21 patients with mitochondrial myopathy, 12 healthy controls and 12 McArdle patients (9). The **Sepsis cohort** was previously described and consists of 98 subjects in four categories: ambulatory controls (n=19), cardiogenic shock (n=19), bacteremia without sepsis (n=18), and septic shock (n=42)(13). The **LSFC cohort** was described previously and consists of 9 patients and 9 controls (11). For the Discovery, Validation, Exercise and LSFC cohorts, informed consent obtained at the time of sample collection did not include provisions for public sharing of metabolomics data; accordingly, raw metabolomics data will not be deposited in public repositories. Raw data for the Sepsis cohort has been deposited at Metabolomics Workbench (Study ID: ST003136).

### Sample collection

For metabolomics, blood samples (collected in either EDTA, sodium heparin or sodium citrate tubes) were centrifuged for 10 min at 2000 rcf at 4°C. Plasma was stored in 500ul aliquots at - 80°C until the time of processing. Urine samples were centrifuged for 10 min at 500 rcf, transferred to a 15ml conical tubes and stored at -80°C until the time of processing.

### Mining existing metabolomics datasets

Metabolomics results from the previously published MELAS **Discovery and Validation cohorts**, the **Exercise cohort**, and the **Sepsis cohort** (see above) were performed by using retention time and exact mass matching from existing datasets run with the same BEH Amide Method (see below). Peak areas were determined for identified unknown metabolites for their relative quantitation.

### LC-MS data collection

**Untargeted metabolomics** was performed on plasma (**LSFC cohort**) and urine (**Discovery cohort**) samples following a previously described method (3). Briefly, we measured polar metabolites with an amide column LC-MS method. 30μL of each plasma sample was extracted with 117μl acetonitrile (Thermo Fisher Scientific) and 20μL of LC-MS grade water (Thermo Fisher Scientific). Samples were vortexed, chilled on ice for 20 min and centrifuged at 21,000xg for 20 min. 100μl of each sample was transferred to an LC-MS vial for analysis. Samples were analyzed using a Q Exactive Plus Orbitrap Mass Spectrometer with a Dionex UltiMate 3000 UHPLC system (Thermo Fisher Scientific). An Xbridge amide HILIC column (2.1 X100 mm, 2.5 μM particle size, Waters) was used to separate metabolites (column temperature of 27°C) and MS was acquired under polarity switching mode. Mobile phase A was 20mM ammonium acetate, 0.25 % ammonium hydroxide, 5% acetonitrile, pH 9.0. Mobile phase B was 100% acetonitrile. The gradient was: 85%B for 0-0.5 minutes, ramp to 35% B from 0.5-9 minutes, ramp to 2% B from 9-11 minutes, hold at 2% B from 11-12 minutes, ramp to 85% B from 12-13.5 minutes, hold at 85% B from 13.5-18 minutes. The flow rate was 0.22 mL/min from 0-14.6 minutes and 0.42 mL/min from 15-18 minutes (and was ramped between 14.6 and 15 minutes). The MS data acquisition was full scan mode in a range of 70–1000 m/z, with resolution 70,000, AGC target of 3E6, and maximum injection time of 80 msec. All LC-MS data were collected with samples injected in a randomized order with pooled QC samples interspersed throughout the runs to quantify instrument performance during the run.

### LC-MS data analysis

Raw untargeted metabolomics profiling data was processed using Progenesis QI (Nonlinear Dynamics; Newcastle upon Tyne, UK) to perform alignment, peak picking, deconvolution and peak identification (Figure 3A, D-F). An in-house retention time library of 520 compounds from IROA Technologies (Bolton, MA) was used for feature identification based on a mass and retention time tolerances of 5ppm and 0.3min, respectively. The Human Metabolome Database compound library was downloaded and searched within Progenesis QI for potential compound matches using a 5ppm mass tolerance (40).

### Absolute quantitation by stable-isotope dilution LC-MS

The absolute concentrations of urine creatinine in the Discovery cohort and plasma lactate and pyruvate in the Exercise cohort were determined using a previously described method (3) that uses the same amide method described above (Untargeted metabolomics). Briefly, we built a calibration curve for each metabolite by preparing nine different concentrations of the unlabeled metabolite in a surrogate matrix (PBS with 30 g/L human serum albumin). Urine, plasma and calibration curve samples were extracted as described above except that a mix of isotopically labeled internal standards were included in the water for amide extraction. Data processing for concentration determinations was performed in TraceFinder (Thermo Scientific). We used the response ratio (unlabeled/labeled) to build a calibration curve for each metabolite which was then used to derive the absolute concentrations in each urine and plasma sample. We fit the linear calibration curves using a 1/x^2^ weighting, excluding the highest and lowest points on the calibration curve until the remaining points fell within 20% of the expected concentration.

### LC-MS fragmentation

Metabolite fragmentation from LC/MS runs was performed with a Q-Exactive Plus mass spectrometer using higher-energy collisional dissociation (HCD) on quadrupole-isolated precursors using a 1 *m/z* isolation window, and captured with a resolution of 70,000 @ 200 m/z.

### GC-MS Methods

Samples were methoxyaminated and trimethylsialylated as described previously (41). Briefly, methoxyamination was first performed by adding 50 μl of methoxyamine hydrochloride (20 mg/ml in pyrimidine) to the dried samples and incubating at 30°C for 90 min. Then, sialylation was carried out by adding 80 μl of MSTFA (N-methyl-N-(trimethylsilyl)trifluoroacetamide) plus 1% TMCS (2,2,2-trifluoro-N-methyl-N-(trimethylsilyl)-acetamide, chlorotrimethylsilane) and incubating at 70°C for 60 min. Derivatized samples were cooled down to room temperature before injection.

GC-MS data was collected on a Q-Exactive GC/MS system (Thermo Scientific) equipped with a TriPlus RSH autosampler, a DB-5MS+DG column (30 m x 0.25 mm x 25 um film, Agilent), and using helium as a carrier gas. The inlet parameters used 1 μL injection of sample and a 10:1 split ratio. The temperature gradient was held at 60°C for 1 min and ramped to 320°C at a rate of 10°C/min, and held at 320°C for 5 min. For electron impact (EI), the MS transfer line and ion source temp were set to 300°C, and the mass spec parameters were a scan range of 40-650 m/z and a resolution of 30,000 at 200 m/z. For chemical ionization (CI), the MS transfer line was set to 300°C and the ion source temperature was set to 200°C, with 1.5 mL/min methane as the CI gas.

### Mouse strains, husbandry, and hypoxia exposure

All animal procedures were conducted in accordance with protocols approved by the Massachusetts General Hospital Institutional Animal Care and Use Committee (IACUC), and animals cared for according to the requirements of the National Research Council’s Guide for the Care and Use of Laboratory Animals. *Ndufs4*^*-/-*^ mice were generously provided by the Palmiter laboratory (University of Washington). Pups were genotyped and weaned at 25-30 d of age. All cages were provided with food and water ad libitum and supporting Napa gel was provided as needed. Mice were maintained in a standard 12h light-dark cycle at a temperature between 20-25°C and humidity between 30% and 70%. *Ndufs4*^*-/-*^ and wild-type (WT) controls were either housed in standard oxygen conditions (21% O_2_) until day 55-60 when *Ndufs4*^*-/-*^ mice present advanced disease symptoms(14) (“normoxia”) or in hypoxia chambers (11% O_2_) starting at day 55 for 1 month which reverses the disease (“hypoxia rescue”), as previously described(15, 16). Male and female mice were used as no sex-specific differences in disease progression have been reported or found in our experience working with these mice. WT and heterozygous mice were used as controls as they are identical in all reported assays performed by us and others. For hypoxia treatments, mice were housed at ambient sea-level pressure in plexiglass chambers, and 11% O_2_ levels were maintained OxyCycler A84XOV Multi-Chamber Dynamic Oxygen Controller (BioSpherix Ltd., Parish, NY) using nitrogen gas (Airgas). The CO_2_ concentration in each chamber, as well as the temperature and humidity, were monitored continuously. Mice were exposed to gas treatment continuously for 24 hours per day, 7 days a week. The chambers were briefly opened three times per week to monitor health status, provide food and water, and change the cages. The Massachusetts General Hospital Institutional Animal Care and Use Committee approved all animal work in this manuscript.

### Mouse tissue preparation for LC-MS

*Ndufs4*^*-/-*^ mice were purchased from The Jackson Laboratory (strain Nr. 027058)(14). *Ndufs4*^*-/-*^ mice with advanced disease and after hypoxia-rescue, and age-matched controls were anesthetized with 5% isoflurane in an induction chamber. Mice were euthanized by terminal blood collection, and which was used for plasma collection using EDTA tube followed by harvesting and clamp-freezing of heart, brain, kidney, liver, and skeletal muscle in liquid nitrogen. Tissues were stored at -80°C until further analysis. Frozen tissues were pulverized into a fine powder with a mortar and pestle at liquid nitrogen/dry ice temperature for metabolite extraction; the powder was transferred into microcentrifuge tubes. ∼10mg tissue powder was transferred to a fresh microcentrifuge tube, and the samples were extracted in 20μL per mg tissue 4:4:2 methanol:acetonitrile:water with 0.1M formic acid, vortexed, and then neutralized with 17.5μL of 150 mg/mL ammonium bicarbonate per 100μL of extraction solution. We processed the mouse plasma as described above (**Untargeted Metabolomics**). Samples from *Tk2*^*Ki/Ki*^ mouse pups along with age-, gender-, and treatment-matched controls (7 untreated *Tk2*^*Ki/Ki*^ mice and 7 matched controls, 9 rapamycin-treated *Tk2*^*Ki/Ki*^ mice and 9 matched controls) were obtained as previously described (22).

### Mouse tissue LC-MS measurements

Processed mouse tissue samples were analyzed for 4,5-DHHA, lactate, and 2-hydroxybutyrate with the BEH amide method as described above (**Untargeted metabolomics**). NADH/NAD^+^ measurements were performed in the presence of stable-isotope labeled internal standard ^13^C_5_ NAD^+^. Samples were then run on the ZIC-pHILIC column (2.1 mm × 150 mm × 5 µm) column held at 40°C at a flow rate of 0.22 mL/min. The gradient was 80% B from 0 - 0.5 mins, ramped to 30% B from 0.5 - 8.5 mins, ramped to 10% B from 8.5 - 8.8 mins, held at 10% B from 8.8 -12.5 mins, ramped to 80% B from 12.5 - 12.8 mins, and held at 80% B from 12.8 - 16 mins. The flow rate was increased to 0.3 mL/min from 13.2 - 15.8 mins. Multiple simultaneous MS SIM scan events were used to improve signal/noise in negative mode at 70,000 resolution (at *m/z* 200): from 1 - 15 mins from 300 *m/z* - 2000 *m/z*, from 4.4 - 5.6 mins 620 *m/z* - 680 *m/z*, from 5.6 - 7.2 mins 720 *m/z* - 780 *m/z*. NADH and NAD^+^ measurements were normalized to the NAD^+^ internal standard, and quantified using a standard curve of unlabeled standards.

### *In vitro* pyruvate dehydrogenase complex measurements

*In vitro* experiments were performed in 250 μL of 136 mM KCl, 10 mM KH_2_PO_4_, pH 7.25 and 2mM MgCl_2_. In the reaction mixture, the final concentrations of pyruvate dehydrogenase (porcine, Sigma) was 0.2 units, with 50 μM thiamine pyrophosphate, 5 mM NAD^+^, 1.5 mM succinic semialdehyde and 1 mM coenzyme A. Finally, the reaction was initiated with the addition of 4 mM pyruvate. For the NADH/NAD^+^ clamping experiment, NAD^+^ concentration was maintained at a constant 5 mM, with 0, 0.5, 2.5, 5, 10, or 15 mM NADH added. At the indicated time point in the time course or 5 minutes for the clamping experiment, each sample was quenched by adding 20 μL water and 117 μL acetonitrile, spun down, and subjected to the same amide method described above in (**Untargeted Metabolomics**). For the NADH/NAD^+^ clamping experiment, the same method was used, but the full scan measurement was performed only in negative ionization mode, and a SIM scan was added from 750-850 *m/z* to improve the measurement of acetyl-CoA.

### Tissue culture experiments

HeLa cells were purchased from ATCC and cultured in DMEM without pyruvate and with 10% dialyzed fetal bovine serum. For measurements, 600,000 cells were seeded per well in a 6-well plate with 2 mL of media and the indicated amount of succinic semialdehyde was freshly added. Cells were treated with 1 μM piericidin or an equivalent amount of DMSO as vehicle control and incubated at either 21% or 1% O_2_. After 24 hours, spent media was collected, and 30 μL media was extracted and subjected to the same amide method described above in (**Untargeted Metabolomics**). For the succinic semialdehyde titration experiment, this method was modified to have a mass spectrometry scan range of 120-1000 *m/z* in negative mode in order to improve measurement sensitivity.

## Acknowledgments

The team is deeply grateful to Dr. Stephen Hersh who, in his role as advisor to the Marriott family and foundation, drove the formation of the unique collaborative initiative that accomplished this work. This work was also enabled by the efforts of Hardik Shah and Drs. Rob Rogers, and Ron Haller who carefully collected samples, analytical data, and phenotypic information in prior studies that were mined for this work. Samples from LSFC cohort and healthy controls were provided through the efforts of many members of the LSFC Consortium (Julie Thompson Legault, John Rioux, Charles Morin, Chantale Aubut, Jeannine Landry, Vanessa Tremblay-Vaillancourt and Jessica Tardif). Catherine Laprise wishes to thank the LSFC patients and their families for their participation in the research. She also acknowledges the Canada research chair and the Fondation du Grand Défi Pierre Lavoie et for their financial support in maintaining the LSFC cohort. This project was supported by the Marriott Mitochondrial Disorders Collaborative Research Network, the Howard Hughes Medical Institute, and the National Institutes of Health (F32GM133047 to OSS, 5R01NS124679-04 to VKM, R01NS117538 to EAS) and the Deutsche Forschungsgemeinschaft (4313887 to MM).

## Supplementary Information

**Fig. S1.**
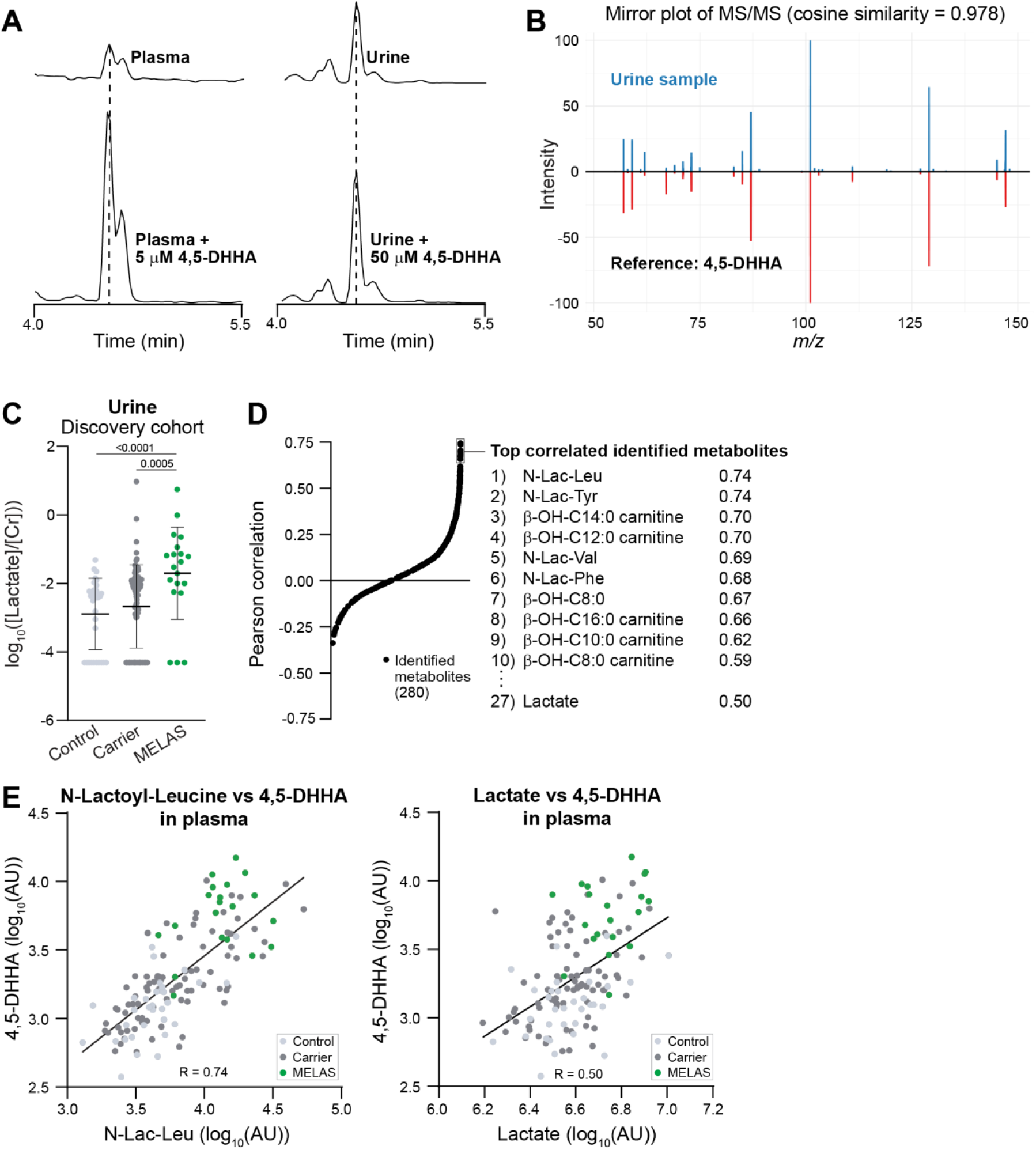
**A)** Extracted ion chromatograms for U147 (147.0662 *m/z*) have the same retention time as each other and as a chemically synthesized pure standard of 4,5-DHHA. **B)** Mirror plot of sample and synthesized 4,5-DHHA shows high cosine similarity score. **C)** Urine lactate/Cr is significantly elevated in MELAS patients compared to Controls and Carriers (lactate concentrations of zero were set to one-tenth the minimum lactate concentration and p-values are based on Mann-Whitney U test). **D-E)** Correlation of 4,5-DHHA with 280 identified metabolites across the Discovery cohort shows it is most strongly correlated with top markers like N-lactoyl-leucine and only moderately correlated with plasma lactate.

**Fig. S2.**
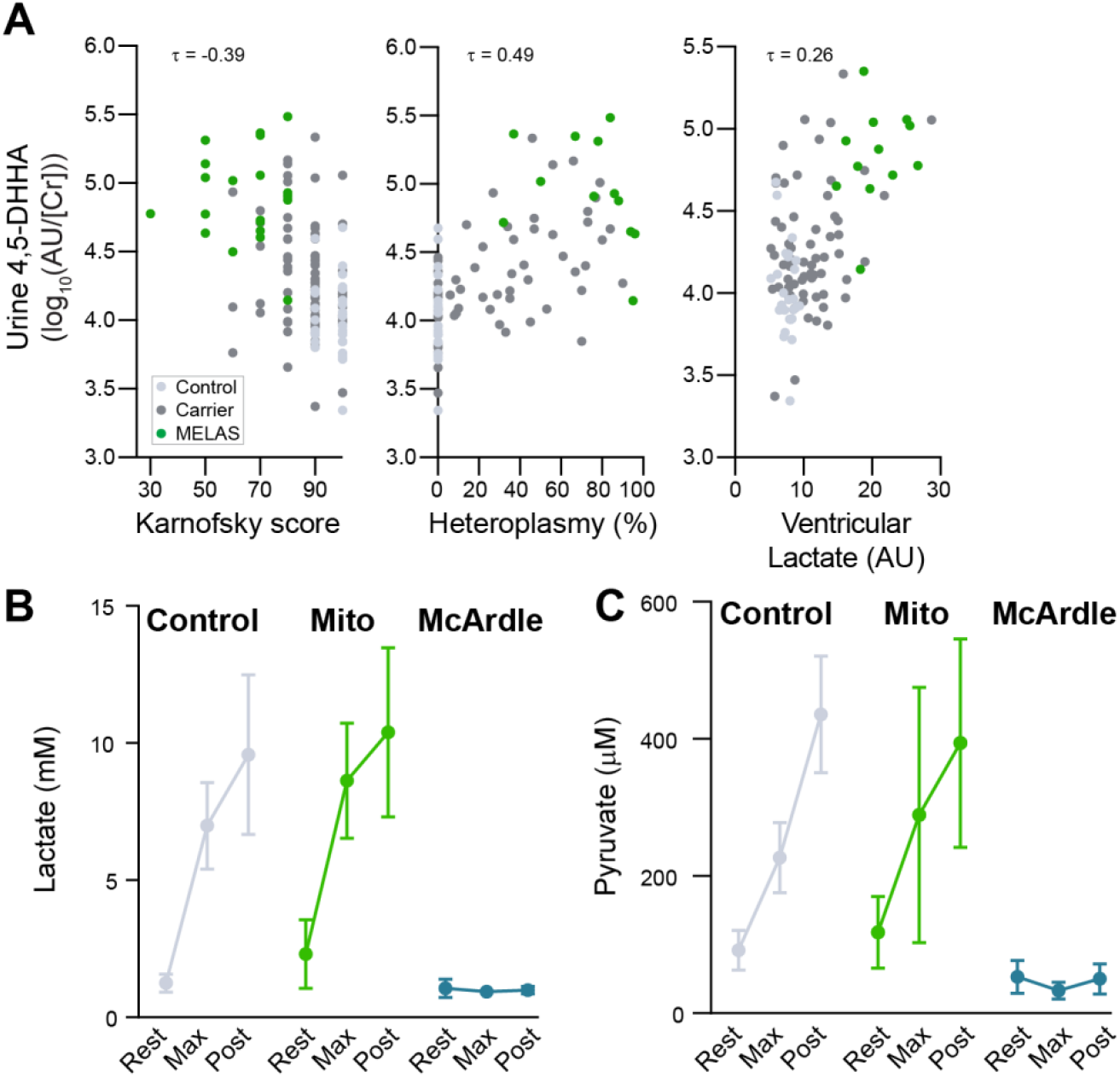
**A)** Urine 4,5-DHHA is strongly correlated with three metrics of disease severity – Karnofsky performance score, urine heteroplasmy, and ventricular lactate. **B, C)** Plasma lactate and pyruvate levels rise during exercise in healthy controls and patients with mitochondrial myopathy while their levels remain low in McArdle disease.

